# UMP inhibition and sequential firing in aspartate transcarbamoylase open ways to regulate plant growth

**DOI:** 10.1101/2020.07.17.208231

**Authors:** Leo Bellin, Francisco del Caño-Ochoa, Adrián Velázquez-Campoy, Torsten Möhlmann, Santiago Ramón-Maiques

## Abstract

Pyrimidine nucleotides are essential to plant development. We proved that Arabidopsis growth can be inhibited or enhanced by down- or upregulating aspartate transcarbamoylase (ATC), the first committed enzyme for *de novo* biosynthesis of pyrimidines in plants. To understand the unique mechanism of feedback inhibition of this enzyme by uridine 5-monophosphate (UMP), we determined the crystal structure of the Arabidopsis ATC trimer free and bound to UMP, demonstrating that the nucleotide binds and blocks the active site. The regulatory mechanism relies on a loop exclusively conserved in plants, and a single-point mutation (F161A) turns ATC insensitive to UMP. Moreover, the structures in complex with a transition-state analog or with carbamoyl phosphate proved a mechanism in plant ATCs for sequential firing of the active sites. The disclosure of the unique regulatory and catalytic properties suggests new strategies to modulate ATC activity and to control *de novo* pyrimidine synthesis and plant growth.

---

Pyrimidine nucleotides are crucial to plant development and growth as components of nucleic acids and as cofactors in the synthesis of sugars, polysaccharides, glycoproteins and phospholipids ^1,2^. However, much remains unknown in plants about the unique organization, regulation, and localization of the enzymes required for *de novo* biosynthesis of pyrimidines, which are potential targets for crop improvement and weed control ^1^. This metabolic pathway starts in the chloroplast, where aspartate transcarbamoylase (ATC) catalyzes the condensation of carbamoyl aspartate from aspartate (Asp) and carbamoyl phosphate (CP) ^1,3^ (Fig. 1a). ATC can be anchored to the inner plastid membrane ^4^ which might facilitate the channeling of carbamoyl aspartate to a cytosolic dihydroorotase (DHO) potentially associated to the outer plastid membrane ^1^. The dihydroorotate produced by DHO diffuses to the mitochondrial intermembrane space and is oxidized to orotate by dihydroorotate dehydrogenase (DHODH), a membrane flavoprotein coupled to the respiratory chain ^3,5^. Orotate returns to the cytosol and is transformed to uridine 5-monophosphate (UMP), the precursor of all pyrimidine nucleotides, in a two-step reaction catalyzed by UMP synthetase (UMPS). Importantly, only in plants, UMP inhibits the activity of ATC, creating a feedback loop that controls the flux through the pathway ^6–9^ (Fig. 1a). UMP also inhibits carbamoyl phosphate synthetase (CPS), the chloroplast enzyme producing CP for both the synthesis of pyrimidines and arginine ^10^, although this effect is counteracted by ornithine to sustain the production of arginine ^11,12^ (Fig. 1a). Thus, ATC is the major regulated enzyme for *de novo* synthesis of UMP in plants ^13^, but yet the feedback mechanism that makes this enzyme different from any other ATC remains uncharacterized.

**Figure 1.**
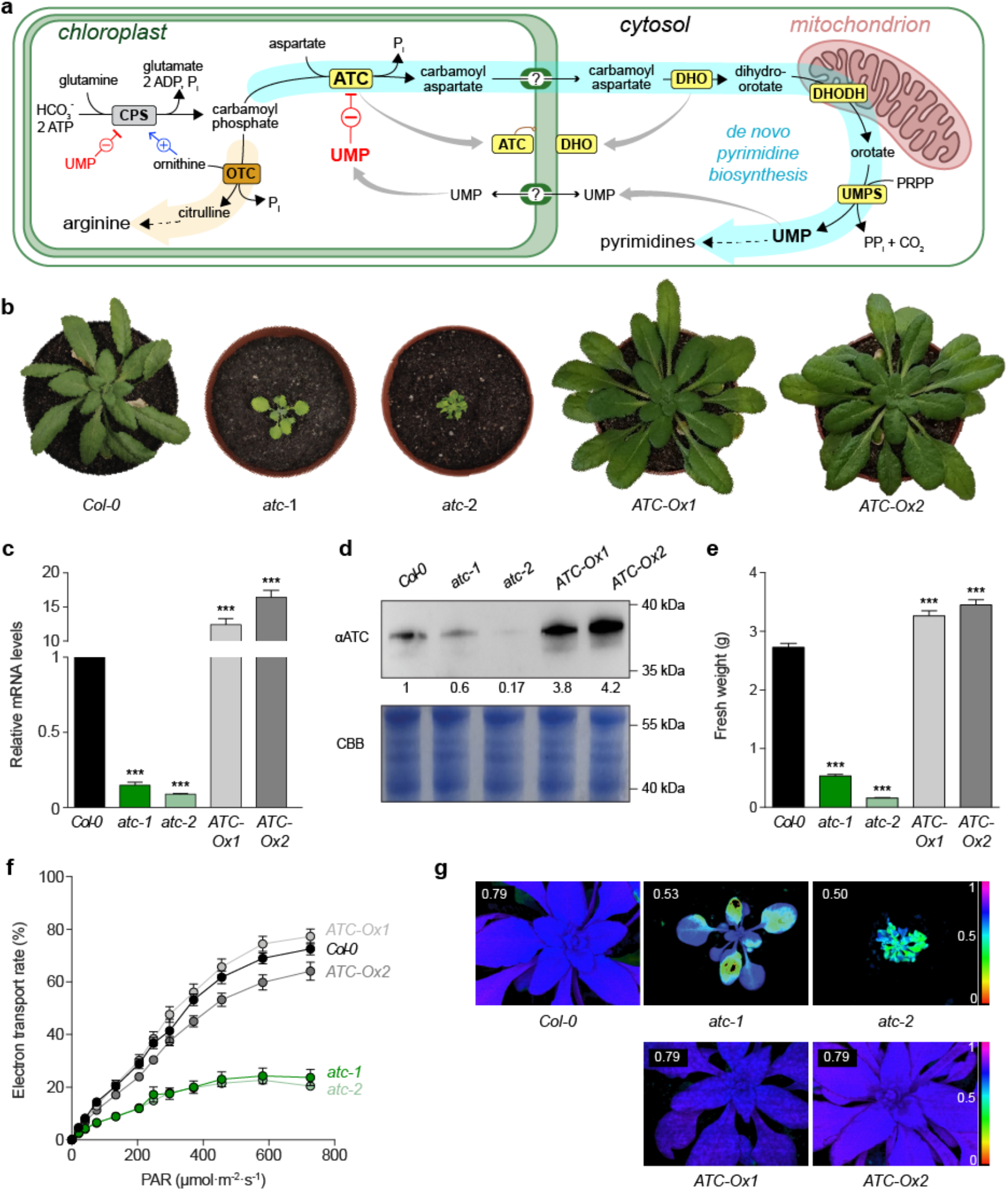
ATC central activity in *de novo* pyrimidine synthesis and plant growth. **a)** Scheme of *de novo* pyrimidine biosynthesis pathway (highlighted in cyan with enzymatic activities in yellow background) and arginine synthesis (in orange) in plants. ATC, aspartate transcarbamoylase; DHO, dihydroorotase; DHODH, dihydroorotate dehydrogenase; UMPS, UMP synthetase; CPS, carbamoyl phosphate synthetase; OTC, ornithine transcarbamoylase. Allosteric inhibition by UMP and activation by ornithine are indicated by red and blue lines, respectively. **b)** Arabidopsis *Col-0* and ATC downregulated (atc-1 and −2) or overexpressing lines (ATC-Ox1 and 2) grown for four weeks in a 14h light/10h dark regime. **c)** *ATC* transcript levels in knockdown and overexpressing lines relative to *Col-0* (n = 3). *ATC* transcript levels relative to actin: 2.2 × 10^−4^ (*Col-0*), 3.48 × 10^−5^ (atc-1), 2.12 × 10^−5^, 2.9 × 10^−4^ (ATC-Ox1) and 3.9 × 10^−4^ (and ATC-Ox2). **d)** Immunoblot with anti-ATC antibody on whole leaf extracts; Coomassie Brilliant Blue (CBB) stained SDS-PAGE was used as loading control. **e)** Fresh weight quantification (n = 10). **f)** Electron transport rate (ETR) determined by PAM in standard light curve setting. **g)** False color presentation of maximal photosynthesis yield monitored by PAM of lines shown in **(f)**. Error bars indicate standard error of the statistical analysis. Asterisks depict significant changes between the different lines referring to *Col-0* control according to one-way ANOVA followed by Tukey’s multiple comparison test (* = p < 0.05, ** = p <0.001, *** = p<0.001)

In general, ATCs consist of a catalytic homotrimer that can be allosterically regulated by association with other proteins. The *Escherichia coli* ATC, for instance, is formed by association of two catalytic trimers with three dimers of regulatory subunits ^14^ responsible for the binding of nucleotides that diminish (UTP and CTP) or enhance (ATP) the activity ^15,16^. Other prokaryotic ATCs lack regulatory subunits and thus, are insensitive to nucleotides ^17–19^. In eukaryotes other than plants, on the other hand, ATC is fused together with CPS into a single multienzymatic protein named CAD that also contains an active DHO (animals) or an inactive DHO-like domain (fungi) ^10,20–24^. This ATC also forms homotrimers, favoring the assembly of CAD into large hexameric particles ^25–28^, and its activity is modulated by the binding of UTP to an allosteric region within CPS ^29,30^. In striking contrast, plant ATCs consist of a UMP-inhibitable homotrimer with no associated subunits, meaning that both the catalytic and regulatory sites must reside within the same polypeptide chain ^31^. However, despite the wealth of biochemical and structural knowledge on ATCs from prokaryotes, fungi and animals, there is no structural information of any plant ATC so far. Thus, the putative binding site for UMP and the catalytic and regulatory mechanisms of ATC in plants remain unknown.

Here we show that the development of *Arabidopsis thaliana* (Arabidopsis) can be severely impaired or enhanced by down- or upregulation of ATC expression. To understand this fundamental activity, we determined the crystal structure of Arabidopsis ATC free and bound to UMP, in complex with a transition-state analog, with CP or bearing a site-specific mutation that turns the enzyme insensitive to UMP. The structural, mutagenesis, and biochemical analyses, disclose unique catalytic and regulatory properties of plant ATCs, suggesting new strategies to control *de novo* pyrimidine synthesis and plant growth.

## RESULTS

### ATC is key for plant development and photosynthetic efficiency

To explore the importance of ATC for plant growth we used artificial microRNA (amiRNA) to knockdown *ATC* in Arabidopsis. Two selected lines, *atc-1* and *atc-2*, exhibited 16% and 10% residual *ATC* transcript and a similar drop in protein levels compared to wild-type (WT; *Col-0*) controls (Fig. 1b-d). Conversely, we constitutively overexpressed *ATC* in two Arabidopsis lines, *ATC-OX1* and *ATC-OX2,* which showed 13- and 18-fold increase in *ATC* transcript and protein levels (Fig. 1b-d). After four weeks on soil, *atc-1 and atc-2* downregulated lines showed a strong reduction of growth, with fresh weights of 19% (0.53 ± 0.09 g plant^−1^) and 6% (0.16 ± 0.03 g plant^−1^) of the *Col-0* control plants (2.73 ± 0.21 g plant^−1^) (Fig. 1b,e). In contrast, *ATC-OX1* and ATC-OX*2* showed increased growth with fresh weights of 119% (3.26 ± 0.09 g plant^−1^) and 126% (3.45 ± 0.09 g plant^−1^) compared to *Col-0*, respectively (Fig. 1b,e).

ATC downregulated lines also showed pale leaves, suggesting lower chlorophyll levels, and presumably impaired photosynthesis, whereas ATC-OX lines exhibited no phenotypic alterations other than the bigger size (Fig. 1b). Because of the pale leaf coloration, four-week-old *atc* plants were subjected to pulse-amplitude-modulation (PAM) fluorometry, which measures chlorophyll fluorescence as an indicator of the photosynthetic capacity. The electron transport rate (ETR) of *atc* lines was less than 20% of *Col-0*, whereas ATC-OX lines showed no significant differences (Fig. 1f). The pronounced ETR decrease was in line with a reduction in the maximal photosynthetic efficiency (Fv/Fm), with values of 0.53 ± 0.03 and 0.50 ± 0.02 for *atc-1* and *atc-2*, respectively, which are markedly lower than the 0.79 ± 0.01 measured in controls (Fig. 1g). Again, Fv/Fm values in ATC-OX lines were similar to *Col-0*.

These results, together with previous studies ^13,32^, demonstrate a regulatory role of pyrimidine *de novo* synthesis in plant growth and a key function of ATC herein.

### Crystal structure of Arabidopsis ATC bound to UMP

To investigate this central enzymatic activity we attempted to produce the ATC from Arabidopsis, a 390 amino acid (aa) precursor protein with an N-terminal chloroplast transit peptide (Fig. 2a and Supplementary Fig. 1). Having difficulties to express the full-length protein in *E. coli*, we tested different N-terminal truncated forms. One construct spanning aa 82–390 (named atATC) was purified as a stable homotrimer (Fig 2b), and matched in size (excluding the 2.6 kDa fusion tag) the 36 kDa mature enzyme in whole leaf extracts (Fig. 2c). atATC produced diffraction quality crystals readily and the structure was determined at 1.7 Å resolution (Table 1; Supplementary Fig. 2). The structure of the atATC trimer resembles a three-bladed propeller with a concave face holding three active sites in between subunits, thus preserving the overall architecture of the transcarbamoylase family ^33^ (Fig. 2d). Each subunit folds into two subdomains of similar size: an N-domain (aa 82–221 and 374–386) occupying the center of the trimer and holding the binding site for CP, and a C-domain (aa 222–373) bearing the Asp binding site (Fig. 2a,d and Supplementary Figs. 1 and 3). The relative orientation of the domains is similar to the open conformation observed in other ATCs crystallized without ligands (Supplementary Fig. 3) ^26,28,34^. In addition, atATC has the CP-loop (aa 156–169) and Asp-loop (aa 309–332) (Fig. 2a,d) that in other ATCs undergo large conformational movements upon substrate binding ^16,28^.

**Figure 2.**
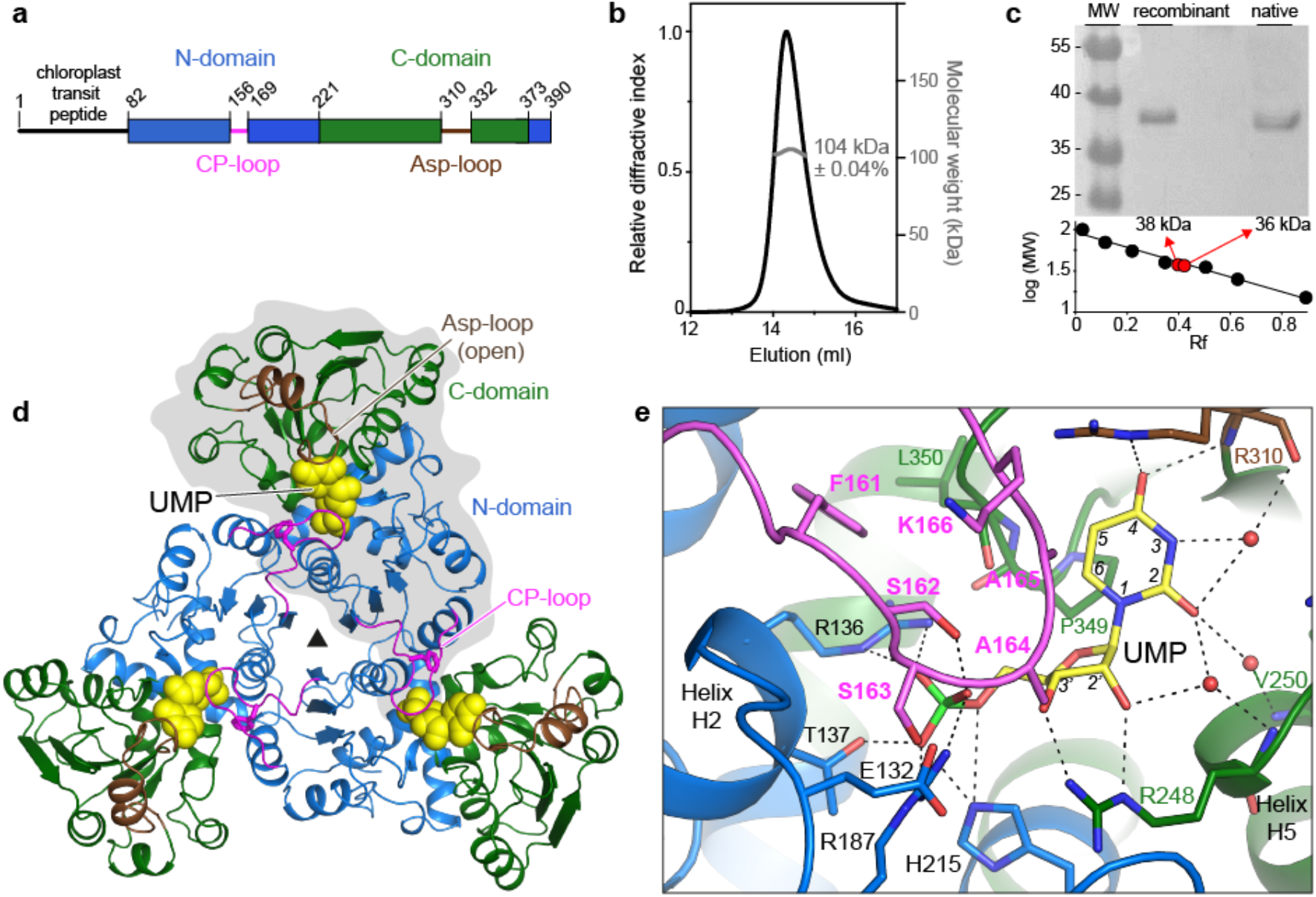
Structure of Arabidopsis ATC bound to UMP. **a)** Scheme of Arabidopsis ATC protein. **b)** SEC-MALS analysis of purified atATC proves the formation of a homotrimer. **c)** Immunoblot of atATC and mature ATC from leaf extract. **d)** Crystal structure of atATC trimer with each subunit bound to one molecule of UMP (shown as yellow spheres). One subunit is shown on grey background. Protein domains are colored as in **a)**. **e)** Detail of the active site with UMP bound. Water molecules are shown as red spheres. Electrostatic interactions are indicated as dashed lines.

**Table 1.** Data collection and refinement statistics.

|  | + UMP | APO | + PALA | + PALA + CP | + CP | F161A + UMP | F161A + PALA |
| --- | --- | --- | --- | --- | --- | --- | --- |
| PDB ID | 6YPO | 6YY1 | 6YS6 | 6YSP | 6YVB | 6YWJ | 6YW9 |
| **Data collection** |  |  |  |  |  |  |  |
| Synchrotron | ALBA (XALOC) | ESRF (MASSIF) | ESRF (MASSIF) | ALBA (XALOC) | ALBA (XALOC) | ALBA (XALOC) | ALBA (XALOC) |
| Wavelength (Å) | 0.9793 | 0.96600 | 0.96600 | 0.97924 | 0.97910 | 0.97926 | 0.97926 |
| Space group | P6<sub>3</sub> | P2<sub>1</sub>2<sub>1</sub>2<sub>1</sub> | P2<sub>1</sub>2<sub>1</sub>2<sub>1</sub> | P2<sub>1</sub>2<sub>1</sub>2<sub>1</sub> | P2<sub>1</sub>2<sub>1</sub>2<sub>1</sub> | P6<sub>3</sub> | P2<sub>1</sub>2<sub>1</sub>2<sub>1</sub> |
| Unit cell a,b,c (Å) | 104.5 104.5 127.8 | 103.7 109.6 208.6 | 86.7 94.9 132.1 | 86.5 94.6 131.8 | 103.3 109.5 212.0 | 104.3 104.3 127.9 | 80.3 98.3 138.8 |
| Resolution (Å) | 52.27 - 1.67<br>(1.73 - 1.67)<sup>a</sup> | 92.85 - 3.07<br>(3.18 - 3.07) | 66.04 - 1.55<br>(1.60 - 1.55) | 76.84 - 1.38<br>(1.43 - 1.38) | 76.16 - 1.83<br>(1.93 - 1.83) | 45.18 - 2.40<br>(2.49 - 2.40) | 46.32 - 1.68<br>(1.71 - 1.68) |
| Unique reflections | 91,002 (9,119)<sup>a</sup> | 45,073 (4,435) | 155,447 (15,241) | 221,343 (21,917) | 212,623 (30,789) | 30,917 (3,526) | 125,454 (6,151) |
| Multiplicity | 10.3 (10.0)<sup>a</sup> | 5.2 (5.2) | 4.8 (5.1) | 6.7 (6.6) | 6.7 (6.9) | 10.0 (10.5) | 6.6 (6.5) |
| Completeness (%) | 99.38 (99.99)<sup>a</sup> | 99.29 (99.57) | 97.4 (96.7) | 99.97 (99.98) | 100 (100) | 100 (100) | 100 (100) |
| R<sub>merge</sub> | 0.1046 (1.396)<sup>a</sup> | 0.14 (0.83) | 0.058 (0.619) | 0.1 (1.154) | 0.051 (1.058) | 0.194 (1.255) | 0.053 (0.840) |
| Mean I/sigma(I) | 12.74 (1.64)<sup>a</sup> | 11.16 (2.33) | 15.07 (2.42) | 10.71 (1.35) | 17.6 (1.7) | 8.5 (2.0) | 18.4 (2.0) |
| Wilson B factor (Å<sup>2</sup>) | 28.26 | 66.93 | 18.98 | 17.49 | 32.00 | 42.10 | 17.94 |
| CC1/2 | 0.997 (0.601)<sup>a</sup> | 0.994 (0.686) | 0.999 (0.752) | 0.994 (0.512) | 0.999 (0.704) | 0.992 (0.629) | 0.999 (0.701) |
| **Model refinement** |  |  |  |  |  |  |  |
| Reflections used in refinement | 91,002 | 45,063 | 155,436 | 221,287 | 201,713 | 29,351 | 125,363 |
| Reflections used for R-free | 4,637 | 2,241 | 7,671 | 11,050 | 10,800 | 1,539 | 6,278 |
| R-factor (%) | 0.15 | 0.19 | 0.12 | 0.14 | 0.17 | 0.14 | 0.15 |
| R-free (%) | 0.17 | 0.25 | 0.15 | 0.17 | 0.21 | 0.19 | 0.17 |
| Average B ($\text{\AA}^2$) | 24.1 | 56.69 | 25.08 | 23.50 | 38.26 | 42.87 | 27.89 |
| Number of non-hydrogen atoms | 5,614 | 14,521 | 8,584 | 8,290 | 17,168 | 4,948 | 8,439 |
| macromolecules | 5,027 | 14,466 | 7,658 | 7,520 | 15,430 | 4,894 | 7,617 |
| ligands | 24 | 12 | 79 | 78 | 214 | 46 | 54 |
| solvent | 563 | 43 | 847 | 692 | 1,524 | 8 | 768 |
| RMS (bonds) | 4 | 6 | 8 | 0.01 | 0.010 | 0.011 | 0.005 |
| RMS (angles) | 0.66 | 0.72 | 0.99 | 1.24 | 1.676 | 1.782 | 0.857 |
| Ramachandran favored (%) | 97.90 | 96.64 | 97.17 | 97.31 | 96.86 | 96.45 | 97.10 |
| allowed (%) | 2.10 | 3.25 | 2.83 | 2.70 | 3.09 | 3.55 | 2.90 |
| disallowed (%) | 0.00 | 0.11 | 0.00 | 0.00 | 0.05 | 0.00 | 0.00 |

<sup>a</sup>Values in between parenthesis correspond to highest resolution shell

Unexpectedly, additional electron density at the active site proved the presence of a molecule of UMP captured during protein expression and kept throughout the purification and crystallization process (Fig. 2d,e and Supplementary Fig. 1). The nucleotide fills the active site, with the ribose in C3’ endo pucker and the base in *anti* conformation (Fig. 2e). The phosphate binds near the N-end of helix H2 and interacts with R136, T137, R187 and H215 (at the N-domain), whereas the ribose 2’- and 3’-OH bind to the side chain of R248 (C-domain). The 4-O atom of the pyrimidine ring interacts with R310 (Asp-loop), and the 2-O and 3-NH bind through three waters to R310, R248 and V250 (C-domain), whereas the C5 and C6 atoms make Van der Waals contacts with the ^349^PLP^351^ loop (C-domain). In addition, the CP-loop from the adjacent subunit interacts with the inhibitor through Van der Waals contacts of residues A164 and A165 and makes a H-bond between S162 and the phosphate (Fig. 2d,e).

Next, we freed the enzyme of UMP by a gel filtration procedure (Supplementary Fig. 4) and determined the crystal structure of the apo form at 3.1 Å resolution (Table 1). The structure turned to be similar to the UMP-inhibited conformation except for aa 160-166 of the CP-loop that appear flexibly disordered in absence of the nucleotide (Supplementary Fig. 3).

### atATC only binds one molecule of PALA per trimer

We investigated the effect of PALA [*N*-(phosphonacetyl)-L-aspartate], a potent ATC inhibitor with structural features of both substrates that mimics the transition-state of the reaction ^35–37^. Seedling assays in presence of 0.2 mM or 0.4 mM PALA showed respectively a decrease in fresh weight to 59% or 23% of untreated seeds and a root length reduction to 29% or 6% (Fig. 3a). These results support the reduced growth observed in *atc* downregulated lines (Fig. 1b-d) and agree with previous PALA-inhibition studies ^36^. Chlorosis was also apparent in PALA-treated seedlings (Fig. 3a), further endorsing the effect of reduced ATC levels on chloroplast functionality (Fig. 1f,g).

**Figure 3.**
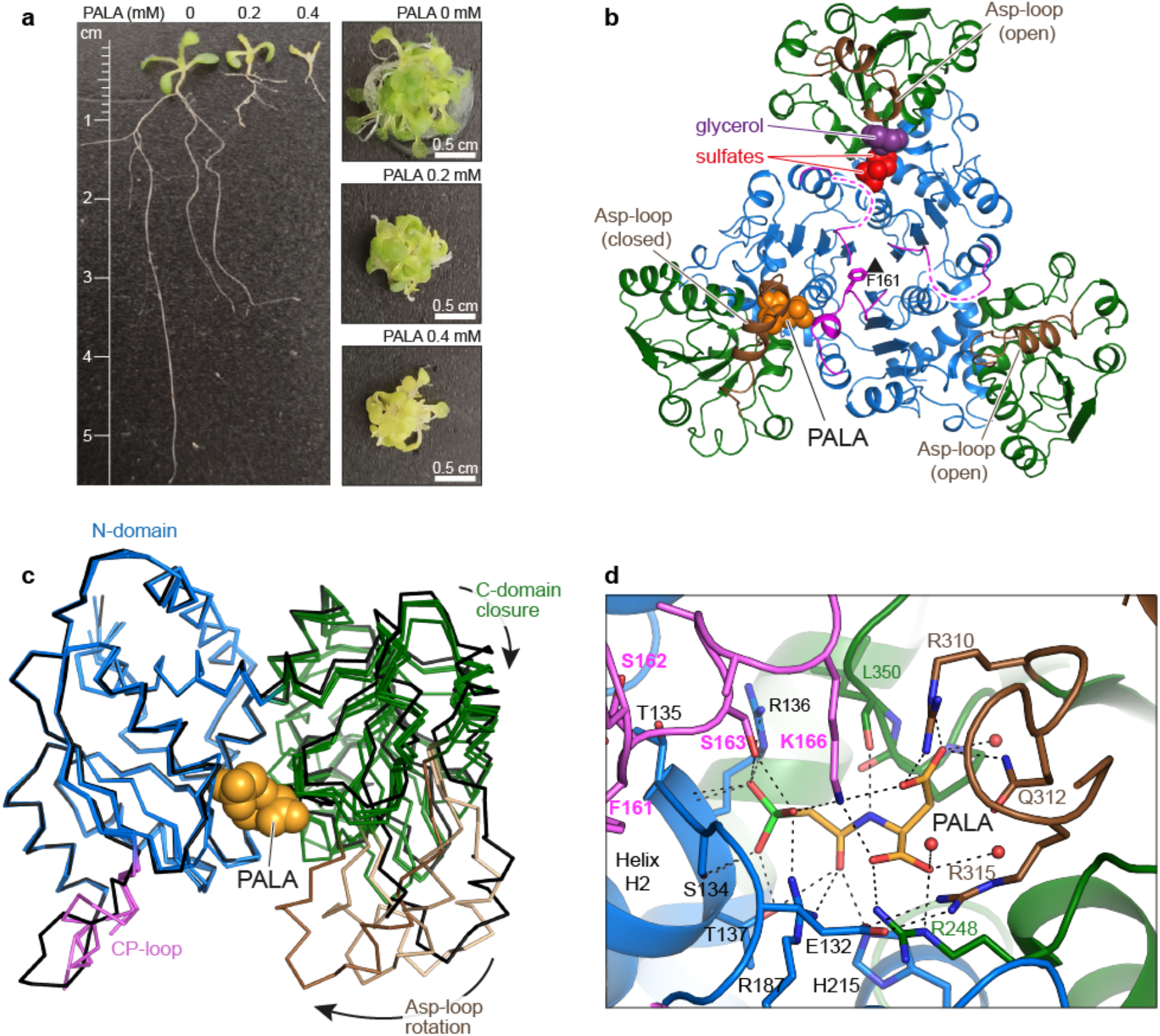
Inhibition of atATC by the transition-state analog PALA. **a)** Inhibition of Arabidopsis seedlings after 7 days treatment with PALA. **b)** Crystal structure of atATC trimer in complex with only one molecule of PALA (shown as orange spheres). Dashed lines indicate flexibly disordered CP-loops. **c)** Superposition of the three subunits in the PALA-bound trimer (colored as in **b)** and in the UMP-bound subunit (in black). Arrows indicate the closure of the subunit upon PALA binding. **d)** Detail of the binding of PALA to the active site. Water molecules are shown as red spheres. Electrostatic interactions are indicated as dashed lines.

To gain further insight into the reaction mechanism, we determined the structure of atATC in complex with the transition-state analog at 1.6 Å resolution (Table 1). Surprisingly, the structure showed the atATC trimer with PALA bound to only one of the subunits (Fig. 3b). This subunit undergoes a 10° hinge closure of the N- and C-domains and a 24° rigid body rotation of the Asp-loop (Fig. 3c), emulating the movement needed to bring CP and Asp in close contact to favor the reaction ^37,38^. In contrast, the other two subunits exhibit an open conformation, similar to the apo or UMP-bound states, and have the active sites empty or with two sulfate ions and one glycerol molecule from the crystallization solution (Fig. 3b). Only the CP-loop interacting with PALA is well-defined in the electron density map (Fig. 3b), whereas the other CP-loops are flexibly disordered.

The substoichiometric binding of PALA is remarkable, since other ATC structures proved the binding of three molecules of PALA per trimer ^28,38–41^ and the interactions with the transition-state analog are virtually identical to those observed in atATC (Fig. 3d and Supplementary Fig. 5). The phosphonate group of PALA binds to the N-end of helix H2 (N-domain) and the O atom of the carbamate moiety interacts with T137, R187 and H215 (N-domain), whereas the N atom binds to L350 (C-domain). Also, the α-carboxylate group binds to R248 (C-domain) and the β-carboxylate binds to R310 and Q312 (Asp-loop). In addition, the CP-loop from the adjacent subunit binds through S163 to the phosphonate moiety, and places K166 at interacting distance of the phosphonate and the α- and β-carboxylates (Fig. 3d).

Isothermal titration calorimetry (ITC) analyses confirmed that PALA binds with high affinity (K_D_^PALA^= 0.6 μM) to only one site per trimer and somehow blocks the entrance of subsequent PALA molecules to the other sites (Table 2). This negative cooperativity effect is specific for PALA, since the unoccupied subunits can still bind CP (K_D_^CP^= 140 μM) or UMP (K_D_^UMP^= 1.2 μM). We also observed negative cooperativity, but to a lesser extent, in the titration with UMP, since the nucleotide binds with high affinity to the first site (K ^UMP^=0.2 μM) and reduces 10-fold the affinity of the other subunits. In turn, CP showed equal affinity for the three active sites (K_D_^CP^ = 77 μM).

**Table 2.** Determination of ligand binding affinities by isothermal titration calorimetry (ITC)

| Sample | Ligand | N<sup>b</sup> | K<sub>D</sub> (μM)<sup>a</sup> |  |  |
| --- | --- | --- | --- | --- | --- |
|  |  |  | Site 1 | Site 2 | Site 3 |
| WT | PALA | 1 | 0.59 | – | – |
|  | UMP | 3 | 0.21 | 2.3 | 1.6 |
|  | CP | 3 | 77 | 77 | 77 |
| WT pre-loaded<br>with PALA | UMP | 2 | *with PALA* | 1.2 | 1.2 |
|  | CP | 2 | *with PALA* | 140 | 140 |
| F161A | PALA | 1 | 0.12 | – | – |
|  | UMP | – | – | – | – |
|  | CP | 3 | 0.73 | 0.73 | 0.73 |

<sup>a</sup> Relative error in K<sub>D</sub> is 30%. When the dissociation constants for a given interaction are equal, ligand binding showed no cooperativity (i.e., independent binding); when the dissociation constants are different, ligand binding showed cooperativity. In all cases, the K<sub>D</sub>'s represent site-specific microscopic dissociation constants for each binding site (i.e., intrinsic site-specific dissociation constant modulated by cooperativity factors if applicable).
<sup>b</sup> Number of binding sites according to the ligand binding model.

These results strongly suggested that despite the overall structural similarity with other ATCs, the atATC trimer hides a mechanism of communication between active sites that allows only one subunit to reach the closed catalytic conformation.

### The CP-loop obstructs the simultaneous closure of the subunits

The explanation for the unusual binding of PALA to atATC was likely at the CP-loop, as the most distinct element compared to other non-plant ATCs (Fig. 2a and Supplementary Fig. 1). This loop is flexible in absence of ligands (Fig. 3b and Supplementary Fig. 3) but adopts two distinct conformations whether UMP or PALA are bound to the adjacent subunit (Figs. 2e and 3d). With UMP, the CP-loop folds in an extended “inhibited” conformation, with A164, A165 and S162 interacting with the nucleotide, S163 and K166 pointing outwards the active site, and the side chain of F161 inserted in between subunits (Fig. 4a). In turn, upon PALA binding, the CP-loop rearranges into two short and nearly perpendicular 3_10_ α-helices, placing S163 and K166 to interact with the transition-state analog and moving A164, A165 and S162 outwards the active site (Fig. 4b). In this “active” conformation, F161 flips 180° compared to the position with UMP, and projects towards the trimer threefold axis, where intersubunit distances are shortened by the interactions between neighbor E156 residues (Fig. 4b). These tight contacts at the center of the trimer are not observed in other ATCs bound to PALA (Supplementary Fig. 5), suggesting that the position of F161 may prevent other CP-loops from reaching a similar active conformation.

**Figure 4.**
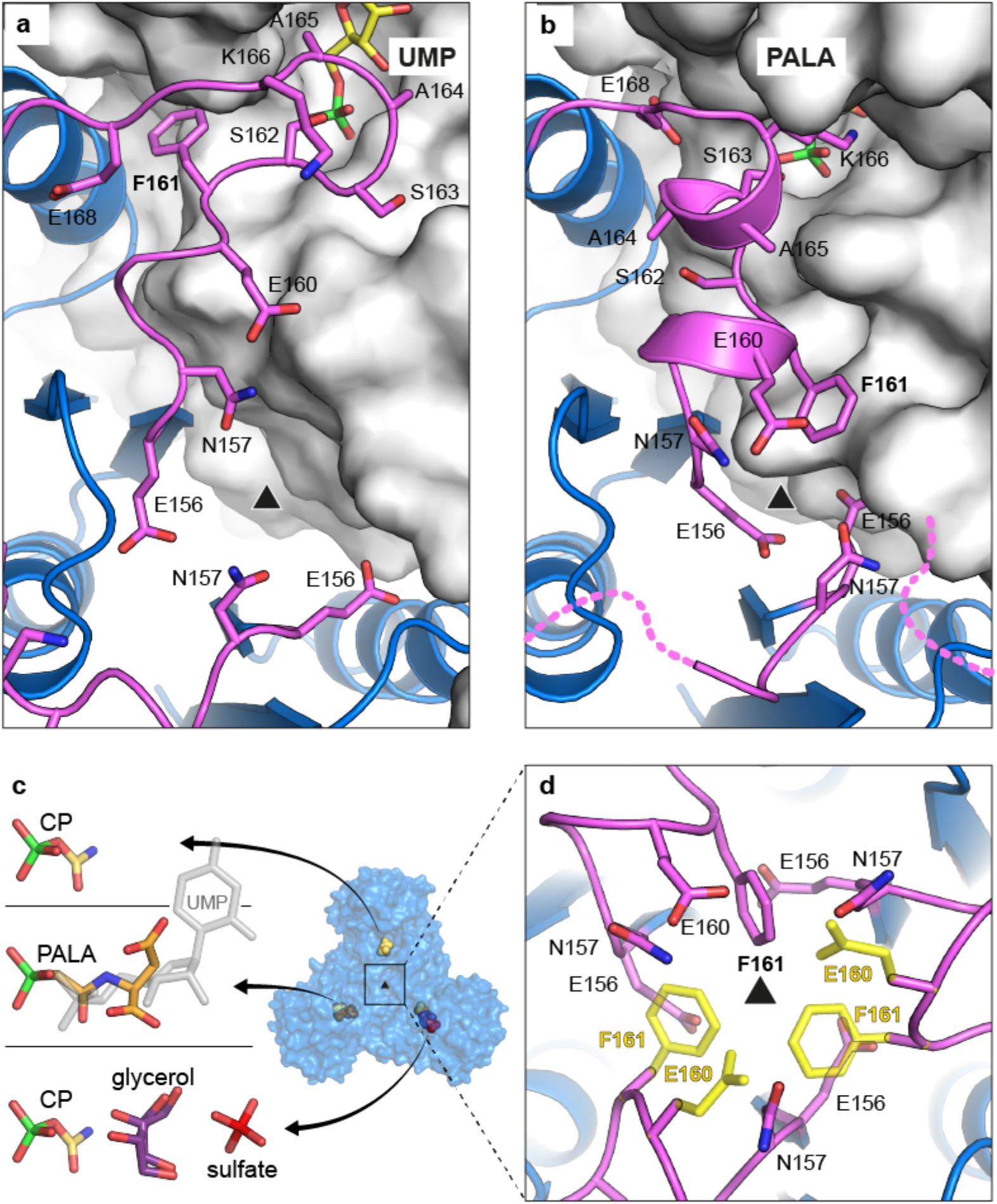
The CP-loop folds in two different conformations for UMP or PALA binding. **a,b)** Cartoon representation of the CP-loop (in magenta) over the active site of the adjacent subunit (shown in surface representation) bound to UMP **(a)** or PALA **(b)**. **c)** Structure of atATC with one subunit bound to PALA, a second one with CP, and the third having CP, glycerol and one sulfate ion. A UMP molecule is shown in semitransparent grey to compare the position relative to the ligands. **d)** Detail view along the threefold axis. The side chains of E160 and F161 are only seen in one subunit. Modelling of these two residues in the other two subunits (shown in yellow semitransparent representation) causes steric clash and charge repulsion.

Two additional atATC structures reinforced this hypothesis. One structure, obtained from crystals with PALA and soaked in CP (Table 1), showed a trimer with one subunit bound to PALA, a second subunit with CP, and a third subunit with CP and with one glycerol and one sulfate ion filling the Asp binding site (Fig. 4c). The second structure, obtained by co-crystallization with CP, showed all three subunits in the trimer bound to CP (Table 1). In both structures, the three CP-loops in the trimer fold in an active conformation but show poor electron density compared to the rest of the protein (Fig. 4d). In fact, E160 and F161 were traced in only one subunit, and modelling in similar conformation in the other subunits caused steric clash and charge repulsion (Fig. 4d).

### Mutation F161A abolishes UMP inhibition and allows binding of three PALA molecules

To further test the role of the CP-loop, we replaced F161 with Ala (F161A). The mutation did not affect the solubility nor the oligomeric state of the protein, but impacted on the enzymatic activity. Initial-rate plots of WT with CP as variable ligand are hyperbolic in absence of UMP (V_max_= 90 nmol·min^−1^·μg^−1^, S_0.5_^CP^= 0.45 mM, S_0.5_^Asp^= 0.94 mM), but turn sigmoidal in presence of UMP, with a Hill-coefficient h= 2.2, indicating positive cooperativity for CP binding (Fig. 5a). In contrast, parallel assays with F161A proved that although the limiting rate is similar to the WT, the activity is not inhibited by UMP and becomes more sensitive to the presence of PALA (Fig. 5a). In addition, F161A showed decreased activity at high substrate concentrations, whereas this substrate inhibition effect was not apparent in the WT (Fig. 5b,c).

**Figure 5.**
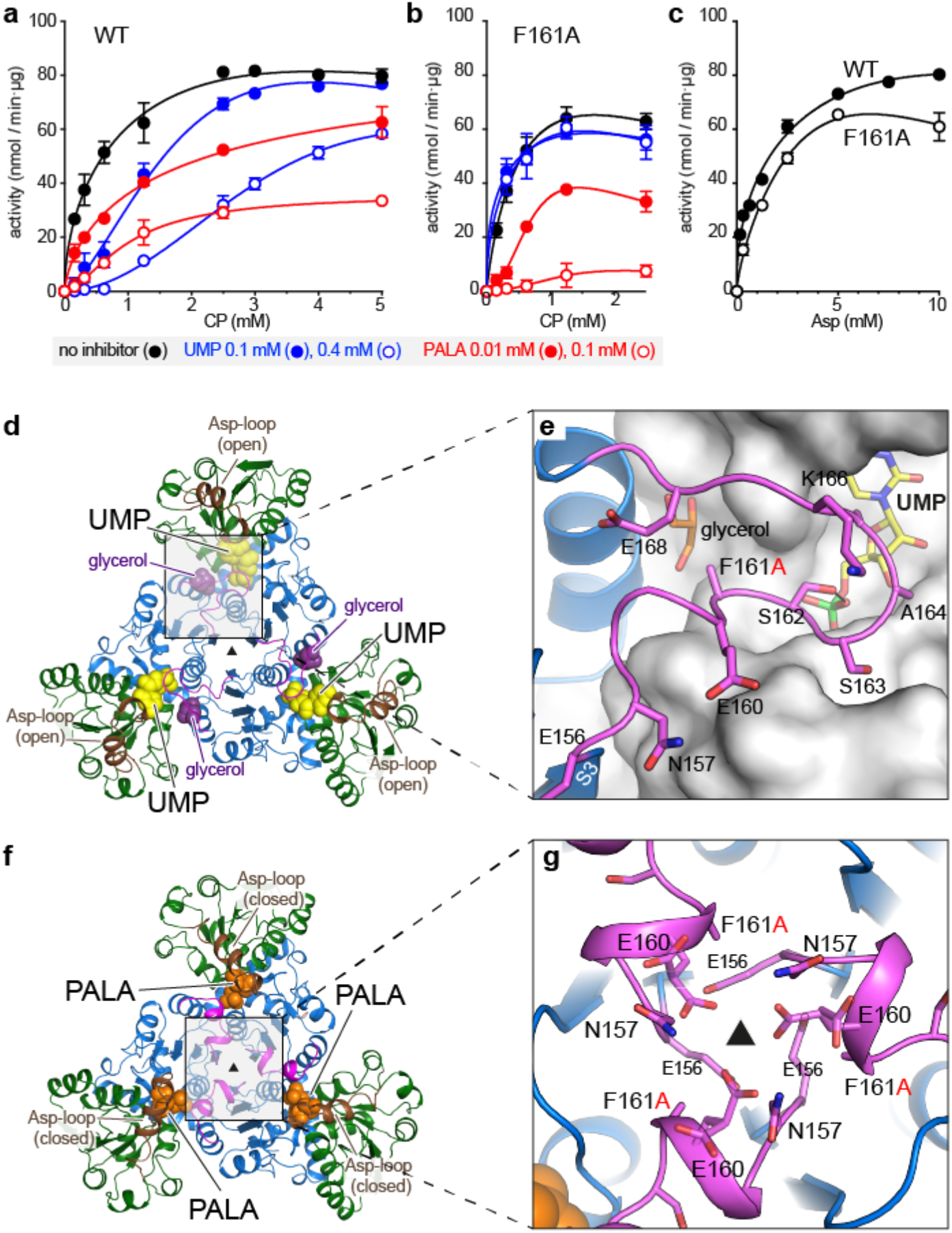
Activity and crystal structure of atATC mutant F161A. **a,b)** CP saturation curves of atATC WT **(a)** and F161A **(b)** in absence and presence of UMP or PALA. **c)** Asp saturation curves for WT and F161A. **d,e)** Crystal structure of atATC-F161A trimer complexed with UMP **(d)**, and detail of the interactions of the CP-loop **(e)**, showing a glycerol molecule replacing the missing F161 side chain. **f,g)** Crystal structure of F161A with PALA bound to the three active sites **(f)**, and detail of the interactions around the molecular axis **(g)**

ITC analysis failed to detect the binding of UMP to F161A, supporting the loss of inhibition by the nucleotide (Table 2). In turn, the affinity for PALA (K_D_^PALA^= 0.12 μM) is increased 5-fold compared to WT, in agreement with the enhanced inhibition, and also the affinity for CP (K_D_^CP^=0.7 μM) is 100-fold higher (Table 2). Likely, removal of the F161 side chain destabilizes the inhibited conformation of the CP-loop, reducing the affinity for UMP, and thus, favoring the alternate CP- or PALA-bound conformation (Fig. 4a,b).

To better understand the effect of the mutation, we determined the structure of F161A with UMP (Table 1). In apparent contradiction with the activity and ITC results, the structure showed a molecule of UMP in the active site (Fig. 5d), likely favored by the high concentration of nucleotide (5 mM) used during crystallization. The CP-loop is in the inhibited conformation, and the missing F161 side chain is replaced by a glycerol molecule (Fig. 5e). We also determined the structure of F161A crystallized with PALA (Table 1). Interestingly, the structure showed a trimer bound to three molecules of PALA rather than one as in the WT (Fig. 5f). Although ITC indicated that PALA binds with high affinity to only one site per trimer (Table 2), the high concentration of PALA (2 mM) in the crystallization condition must favor a low-affinity binding to the other subunits. Importantly, the three CP-loops fold in an active conformation, have well-defined electron density, and show no steric clashes around the molecular threefold axis, since the bulky F161 side chain is missing and E160 adopts alternate conformations (Fig. 5g).

## DISCUSSION

### ATC exerts highest control over *de novo* pyrimidine synthesis

The pathway for *de novo* synthesis of UMP is evolutionary conserved in all plants examined so far, and loss of function of any of the enzymes involved is presumably lethal ^42,43^. However, downregulation of CPS or DHODH had no effect on potato or tobacco plant growth and DHO mutants were only slightly affected ^43^, whereas downregulation of UMPS led to increased yield of potato tuber by overcompensation reactions from the pyrimidine salvage pathway ^44^. In contrast, we showed that ATC downregulation strongly inhibits plant growth (Fig. 1b-e), as previously reported for *Solanaceae* and Arabidopsis plants ^42,43^, causing a severe decrease in photosynthetic efficiency (Fig. 1f,g). Conversely, we proved for the first time that plant growth can be enhanced by ATC overexpression (Fig. 1b-f). Likely, ATC is not produced in large excess ^43^, and thus, plants appear specially sensitive to ATC levels, which exert highest control over pyrimidine *de novo* synthesis. The production of ATC is under transcriptional regulation in response to tissue pyrimidine availability ^42,43^ and to growth signals mediated by the TOR pathway ^45^. However, transcription and translocation of newly synthesized ATC into the chloroplast are slow and energetically costly processes that do not correct for rapid fluctuations needed to maintain nucleotide homeostasis. For this, allosteric regulation by UMP is the major mechanism controlling ATC activity in plants ^7^. However, until know we lacked detailed information of how this feedback loop happened.

### Feedback regulation by UMP

The unprecedented structure of an ATC from plants reveals the mechanism of UMP inhibition. Rather than occupying an allosteric pocket, UMP binds and blocks the active site (Fig. 2), directly competing with CP, the substrate binding in first place ^18,46^. The capacity to bind UMP relies on small changes localized in the CP-loop (Figs. 2e and 4a), since other UMP-interacting elements are common to all ATCs (Supplementary Fig. 1). Indeed, a single point mutation in the CP-loop, F161A, is sufficient to turn atATC insensitive to UMP without affecting the activity nor the inhibition by PALA (Fig. 5 and Table 2). Since the sequence of the CP-loop appears invariant in all plant ATCs known up to date (Supplementary Fig. 1), we propose that the UMP-inhibition mechanism described here for Arabidopsis ATC must be conserved across the plant kingdom. Thus, unlike other ATCs that rely on complex associations with regulatory proteins, the current structures explain how plant ATCs have evolved to maintain a simple organization with both catalytic and regulatory capacities, finding a convenient solution for transcriptional regulation and translocation to the plastid of a single gene product.

The structure also explains the unsolved problem of why plant ATCs are selectively inhibited by UMP and not by any other nucleotide ^8,35^. UMP blocks the subunit in a wide-open conformation (Supplementary Fig. 3), where the N- and C-domains cannot move further apart to accommodate a di- or tri-phosphorylated nucleotide. Also, the pocket for the nitrogenous base is too small for the double ring of a purine and highly selective for uracil, since the methyl group of thymine would clash with the ^349^PLP^351^ loop, whereas the cytidine amino group would distort the interaction with R310 (Fig. 2e). Finally, one would expect the binding of deoxy-UMP to be weak based on the interaction between the ribose OH groups and the side chain of R248, which mimic the recognition of the Asp α-COOH group (Fig. 2e).

atATC has a surprising affinity for UMP, ~400-fold higher than for CP (Table 2), which is explained by the extensive contacts of the nucleotide with the CP and Asp binding sites, similar to what PALA does. However, whereas binding of PALA involves large conformational changes (Fig. 3c), UMP binds to the more energetically favorable open state, and this might explain the 3-fold higher affinity for UMP than for the transition-state analog (Table 2). It is uncertain whether such affinity for the natural inhibitor is retained *in vivo* or if it could be further increased by interaction with phospholipids, as reported for wheat germ ATC (Khan et al., 1996). In any case, based on these results one would expect that ATC is constitutively repressed under steady state conditions where UMP is estimated to reach sub-millimolar concentrations ^43,44,47,48^. However, other features found in atATC suggest that this important activity is fine-tuned by the balance of UMP and CP contents in the cell. Indeed, we showed that the affinity for UMP is modulated by the communication between subunits, so that activity is diminished by high-affinity binding of the inhibitor to one subunit, whereas complete inactivation requires a 10-fold increase in the UMP pool (Table 2). On the other hand, the response to UMP also varies with the concentration of CP, as shown by the change in the kinetic curves from a hyperbola to a sigmoid (Fig. 5a). This behavior, which was first described for wheat germ ATC ^47^, could be relevant for the coordination of *de novo* pyrimidine and arginine synthesis, two pathways that depend on CP availability ^11,12^ (Fig. 1a). For instance, when pyrimidine pools are low, the activity of UMP-free ATC responds linearly with the concentration of CP (Fig. 5a, hyperbola), competing with arginine synthesis to ensure the production of nucleotides. Then, as UMP pool builds up, the synthesis of pyrimidines is reduced by the end product inhibition of ATC and CPS. However, upon CPS inhibition, ornithine accumulates and reverses the UMP inhibition on CPS, providing CP for arginine synthesis. Meanwhile, ATC activity remains negligible at low CP concentrations (Fig. 5a, sigmoid), but would sharply increase if the CP pool allows feeding both metabolic pathways.

### Sequential firing of active sites

atATC provided the unprecedented example of an ATC binding to a single molecule of PALA per trimer (Fig. 3b and Table 2). Since PALA is a transition-state analog, this finding indicates that catalysis in plant ATCs might occur only in one subunit of the trimer at a time. This unexpected mechanism relies on the projection of F161 towards the threefold axis, which prevents the CP-loops from reaching simultaneously the active conformation (Fig. 4b,d). Thus, F161 plays a dual role both stabilizing the UMP inhibited conformation and synchronizing the firing of the subunits in the trimer. Indeed, mutation F161A does not only turn the enzyme insensitive to UMP but also allows the binding of three PALA molecules per trimer (Fig. 5e), although only one does it with high affinity (Table 2), suggesting that other elements might contribute to the communication between subunits. On the other hand, it is tempting to speculate that the activation of the subunits could follow a specific order. Certainly, the atATC structure with PALA and CP provides a suggestive snapshot of each subunit at a different stage of the reaction (Fig. 4c), and further studies should explore this possibility.

The alternation in the firing of the active sites might not be exclusive of plant ATCs. Indeed, human and *E. coli* ATCs exhibit much lower affinities for the binding of PALA to the third active site ^28,49^, and this led us to suggest that only two subunits per trimer were catalytically competent at a time ^28^. The current results further stress the idea that ATCs in general might work more efficiently if not all subunits react simultaneously, avoiding intersubunit contacts that could slow down the conformational movements needed during catalysis ^28,50^ (Fig. 3c). Indeed, the obstruction between subunits would explain the decreased activity at high substrate concentrations, a well-characterized but poorly understood phenomenon in ATCs ^50^. Interestingly, atATC does not exhibit substrate inhibition (Fig. 4d), in agreement with the existence of a mechanism to prevent the closure of more than one subunit at a time. In turn, F161A bypasses this mechanism and shows substrate inhibition (Fig. 5b,c), although less acute than other ATCs ^46,50^, which might explain why the activity is not higher than WT despite having three active sites that could fire simultaneously.

## Conclusions

The *de novo* pyrimidine synthesis pathway is a promising target for biomedical and biotechnological intervention ^51–55^. Our analysis has uncovered the unique regulatory and catalytic mechanisms of plant ATCs, suggesting new strategies to modulate this central enzymatic activity and thus, inhibit or enhance plant growth in a similar manner as we have shown by down- or upregulating ATC expression (Fig. 1b). For instance, the design of new herbicides could be based on UMP analogs with additional groups replacing the observed water-mediated interactions (Fig. 2e), or on compounds targeting the cavity at the threefold axis and blocking the sequential activation of the CP-loops. On the other hand, it should be possible to enhance growth of transgenic crops expressing UMP-insensitive and thus, constitutively active ATC. The present results should guide in the design of highly efficient and non-regulated ATC variants of biotechnological interest.

## ACKNOWLEDGEMENTS

This work was supported by funding from the Spanish Ministry of Science, Innovation and Universities (BFU2016-80570-R and RTI2018-098084-B-100; AEI/FEDER, UE) to SRM, DFG (Mo 1032/4-1 and CRC175-B08) to TM, DFG-IRTG 1830 to TM and LB, and Bayer AG Crop Science (Grants for Targets 2017-02-005) to SR and TM. We thank the staff from ALBA (Barcelona, Spain) and ESRF (Grenoble, France) synchrotron facilities for help during crystallographic data collection; R. Campos-Olivas and C.M. Santiveri for support with SEC-MALS analyses; S. Niopek-Witz and N. Navaseelan for support with ATC expression and kinetic analysis; L. Ohler, A. John and A. Grande-García for support during protein purification, mutant generation and crystallization trials; and M. Moreno-Morcillo for support with crystal structure analysis. We thank D. Lietha and E. Neuhaus for critical reading of the manuscript.

## MATERIALS AND METHODS

### Plant growth

For DNA isolation, tissue collection and phenotypic inspection, wild-type *Col-*0 and transgenic *Arabidopsis thaliana* (L.) Heynh. plants (ecotype Columbia) were used throughout. Plants were grown in standardized ED73 (Einheitserde und Humuswerke Patzer) soil under long day conditions (120 μmol quanta m^−2^ s^−1^ in a 12 h light and 12 h dark regime, temperature 22°C, humidity 60%). Prior to germination, seeds were incubated for 24 h in the dark at 4°C for imbibition ^68^. Alternatively, to assess the effect of PALA on growth, plants were cultivated under short-day conditions in liquid culture according to a protocol suitable for fresh weight determination and feeding of effector molecules ^69^. For growth experiments under sterile conditions the seeds were surface sterilized in 5% sodium hypochloride before adding them to the ½ MS liquid medium supplemented with 1% (w/v) sucrose with or without 0.1 mM, 0.2 mM or 0.4 mM PALA. The liquid cultures were maintained on a shaker at 100 rpm under the same light and temperature conditions as soil grown plants. Leaf extract of wild type and mutants was prepared by homogenizing leaf material in extraction buffer (50 mM HEPES-KOH pH 7.2, 5 mM MgCl_2_, 2 mM phenylmethylsulfonyl fluoride (PMSF) on ice. The homogenous extract was centrifuged at 20,000g for 10 min at 4°C. The supernatant was collected and stored on ice until use.

### Construction of ATCase knock down and overexpressor plants

ATC (*pyrB;* At3g20330) knock down mutants were generated using an established protocol for gene silencing by artificial microRNA (amiRNA) ^70^. An amiRNA targeting *pyrB* was designed using an online tool (http://wmd3.weigelworld.org). The sequence *TAATGACAGGTATATCGGCAG* was used for generation of primers, and Gateway™ compatible sequences attP1 and attP2 were added to primers to engineer the amiRNA fragment (Supplementary Table S1). Subsequently, the fragment was sub-cloned via BP-clonase reaction into the Gateway™ entry vector pDONR/Zeo and via LR-clonase reaction into the destination vector pK2GW7, which contains a 35S-CaMV promotor. Several independent lines were obtained exhibiting 16-10% of transcript reduction and two were selected for further analysis. *ATC* overexpressor plants were generated by cloning full length *ATC* (for primers see Supplementary Table S1) using Gateway technology into pUB-Dest under the control of the ubiquitin-10 promoter ^71^.

### Gene Expression Analyses

Leaf material of soil grown plants was collected and homogenized in liquid nitrogen prior to extraction of RNA with the Nucleospin RNA Plant Kit (Macherey-Nagel, Düren, Germany) according to the manufacturer’s advice. RNA purity and concentration were quantified using a Nanodrop spectrophotometer. Total RNA was transcribed into cDNA using the qScript cDNA Synthesis Kit (Quantabio, USA). qPCR was performed using the quantabio SYBR green quantification kit (Quantabio) on PFX96 system (BioRad, Hercules, CA, USA) using specific primers (Supplementary Table S1), and At2g3760 (Actin) was used as reference gene for transcript normalization.

### Pulse-Amplitude-Modulation (PAM) Fluorometry Measurements

A MINI-IMAGING-PAM fluorometer (Walz Instruments, Effeltrich, Germany) was used for *in vivo* chlorophyll A light curve assays on intact, 6 week-old dark-adapted plants using standard settings ^72^.

### Protein production

Arabidopsis ATC sequence encoding aa 82–390 was PCR amplified from cDNA using a pair of specific primers (Supplementary Table S1) and transferred via NdeI-XhoI to a modified pET28a plasmid ^56^. Mutant F161A was made using the Quick-Change II-E site-directed mutagenesis kit (Stratagene) and a pair of complementary primers (Supplementary Table S1). Transformed *E. coli* BLR(DE3)-pLysS cells (Merck) were grown overnight at 37°C in Terrific Broth (TB) medium containing 25 μg·ml^−1^ kanamycin, 10 μg·ml^−1^ tetracycline and 25 μg·ml^−1^ chloramphenicol to an optical density at 600 nm of 0.7-0.9. Protein expression was induced with 0.5 mM isopropyl β-D-thiogalactopyranoside (IPTG) overnight at 37°C. Cells were resuspended in buffer A (20 mM Tris– HCl pH 8.0, 0.5 M NaCl, 10 mM imidazole, 5% glycerol, 2 mM β-mercaptoethanol) and disrupted by sonication. The clarified lysate was incubated with Ni-sepharose 6 fast flow beads (GE Healthcare) equilibrated in buffer A. After washing with buffer A supplemented with 40 mM imidazole, the protein was eluted in buffer A with 300 mM imidazole. The protein was further purified by size-exclusion chromatography on a Superdex 75 10/300 column (GE Healthcare, USA) equilibrated in buffer GF [20 mM Tris pH 7.0, 0.1 M NaCl, 2% glycerol, 0.2 mM Tris(2-carboxyethyl) phosphine (TCEP)]. The protein was concentrated to 5 mg·ml^−1^ in an Amicon ultracentrifugation device (Millipore) with 10 kDa cutoff and directly used for further studies or supplemented with 40% glycerol, flash-frozen in liquid nitrogen and stored at −80 °C. All purification steps were carried out at 4 °C and the purity of the sample was evaluated by SDS-PAGE and Coomassie staining. Protein concentration was determined by Bradford assay. To remove UMP from the purified protein, the sample was diluted to 1 mg·ml^−1^ and supplemented with 50 mM CP and 100 mM Asp, and filtered at room temperature through three consecutive PD-10 desalting columns (GE Healthcare) equilibrated in GF buffer containing 50 mM CP and 100 mM Asp. In between columns, the 3.5 ml eluted sample was concentrated down to 2.5 ml using an Amicon ultracentrifugation device. In the last step, the sample was filtered through a PD-10 in GF buffer without substrates.

### Size-exclusion chromatography coupled to multi-angle light scattering (SEC-MALS) measurements

For molar mass determination, 400 μl of purified protein at 1.4 mg·ml^−1^ was fractionated on a Superdex 200 10/300 column equilibrated in GF buffer using an AKTA purifier (GE Healthcare) at a flow rate of 0.5 ml·min^−1^. The eluted sample was characterized by in-line measurement of the refractive index and multi-angle light scattering using Optilab T-rEX and DAWN 8+ instruments, respectively (Wyatt). Data were analyzed with ASTRA 6 software (Wyatt) to obtain the molar mass, and plotted with software GraphPad.

### Immunoblotting

15 ng of recombinant ATC or 15 μg of a fresh protein extract from Arabidopsis leaves separated in a 15% SDS-PAGE gel were transferred onto a nitrocellulose membrane (Whatman, Germany) by wet blotting. The membrane was blocked in phosphate-buffered saline plus 0.1% [v/v] Tween 20 (PBS-T) with 3% milk powder for 1 h at room temperature, followed by three washes of 10 min in PBS-T. Then, the membrane was incubated with a rabbit polyclonal antiserum raised against recombinant ATC (Eurogentec, Belgium) for 1 h, followed by three washes with PBS-T. Next, the membrane was incubated for 1 h with a horseradish peroxidase (HPR) conjugated anti-rabbit antibody (Promega, Walldorf, Germany) diluted in PBS-T with 3% milk powder. The result was visualized by chemiluminescence using the ECL Prime Western blotting reagent (GE Healthcare) and a Fusion Solo S6 (Vilber-Lourmat) imager.

### Crystallization, diffraction data collection and structure determination

Initial crystallization screenings were performed at room temperature using the sitting-drop vapor diffusion method and 96-well MRC plates (Hampton). Drops consisting of 0.7 μl protein solution at 5 mg·ml^−1^ plus 0.7 μl reservoir solution were equilibrated against 60 μl of reservoir solution using JCSG+, PACT, and Crystal Screen (Hampton Research) commercial screens. Diamond and rod-shape crystals appeared after few hours. Initial hits were further optimized in 24-well sitting-drop plates (Qiagen) using as reservoir solution 18-22 % PEG 3350, 0.1 M Na_2_SO_4_ and 0.1 M Bis-Tris pH 6.5. The protein freed of UMP was crystallized in 18-22% PEG 3350 and 150– 200 mM potassium acetate. Also, the protein was co-crystallized with 2 mM PALA or 10 mM CP in 25% PEG 3350, 0.2 M Li_2_SO_4_ and 0.1 M Bis-Tris pH 5.5. Crystals of mutant F161A were obtained in similar conditions as the WT protein. All crystals were cryo-protected in mother liquor supplemented with 20% glycerol and flash-frozen in liquid nitrogen. X-Ray diffraction data were collected at BL13-XALOC (ALBA synchrotron, Barcelona) or MASSIF (ESRF, Grenoble) beamlines using Pilatus 6M detectors. Data processing and scaling were performed with XDS ^57^ and autoPROC ^58^. Crystallographic phases were obtained by molecular replacement using PHASER ^59^ and the structure of the ATC domain of human CAD (PDB 5G1O) ^28^ as the search model. The models were constructed by iterative cycles of model building in COOT ^60^ and refinement with PHENIX ^61^ or Refmac5 ^62^ in CCP4 ^63^. Figures were prepared with PyMOL.

### Isothermal Titration Calorimetry

Experiments were performed in an Auto-iTC200 calorimeter (MicroCal, Malvern-Panalytical) at 25° C with a 0.2-ml reaction cell and 30–40 μM protein solution in the cell. Titration experiments consisted of 19 injections of 2 μl of 0.4 mM PALA, 0.4 mM UMP or 1 mM CP. Binding of UMP or CP was also performed on 30–40 μM protein samples pre-incubated with 80 μM PALA. All solutions were in GF buffer and degassed and mixed in the cell by stirring at 750 rpm. Data analysis was performed with Origin 7 (OriginLab) using a general three-site cooperative and non-cooperative binding model, restricted to one or two ligand binding sites when convenient ^28,66,67^. Non-linear least-squares regression analysis was employed to estimate dissociation constants for the interaction of the different ligands with ATC. The number of ligand binding sites or stoichiometry was fixed by the model employed for the analysis (one site, two sites, or three sites); in all experiments the fraction of active or binding-competent protein was close to 1.

### Activity assays

Activity was assayed by a colorimetric method ^64^ adapted to a 96-well plate format ^28,65^. The reaction was carried out in 50 mM Tris-acetate (pH 8.3) and 0.1 mg·ml^−1^ BSA in a final volume of 150 μl. atATC was pre-incubated with Asp for 10 min in a water bath at 25 °C and the reaction was triggered by adding CP and stopped after 5 min with 100 μl of color solution. Reaction tubes were closed, boiled at 95 °C for 15 min, and kept in the dark for 30 min before measuring the absorbance at 450 nm in a Tecan infinite m200 microplate reader (Tecan). Substrate saturation curves were done varying the concentration of one substrate maintaining a fix concentration of the other substrate (5 mM for CP or 10 mM for Asp). Protein concentration in the assay was 0.3 μM (0.01 mg·ml^−1^) both for WT and F161A. Inhibition was measured by adding PALA or UMP during the pre-incubation. Data analysis was done with GraphPad.

## SUPPLEMENTARY INFORMATION

**Supplementary table S1.**
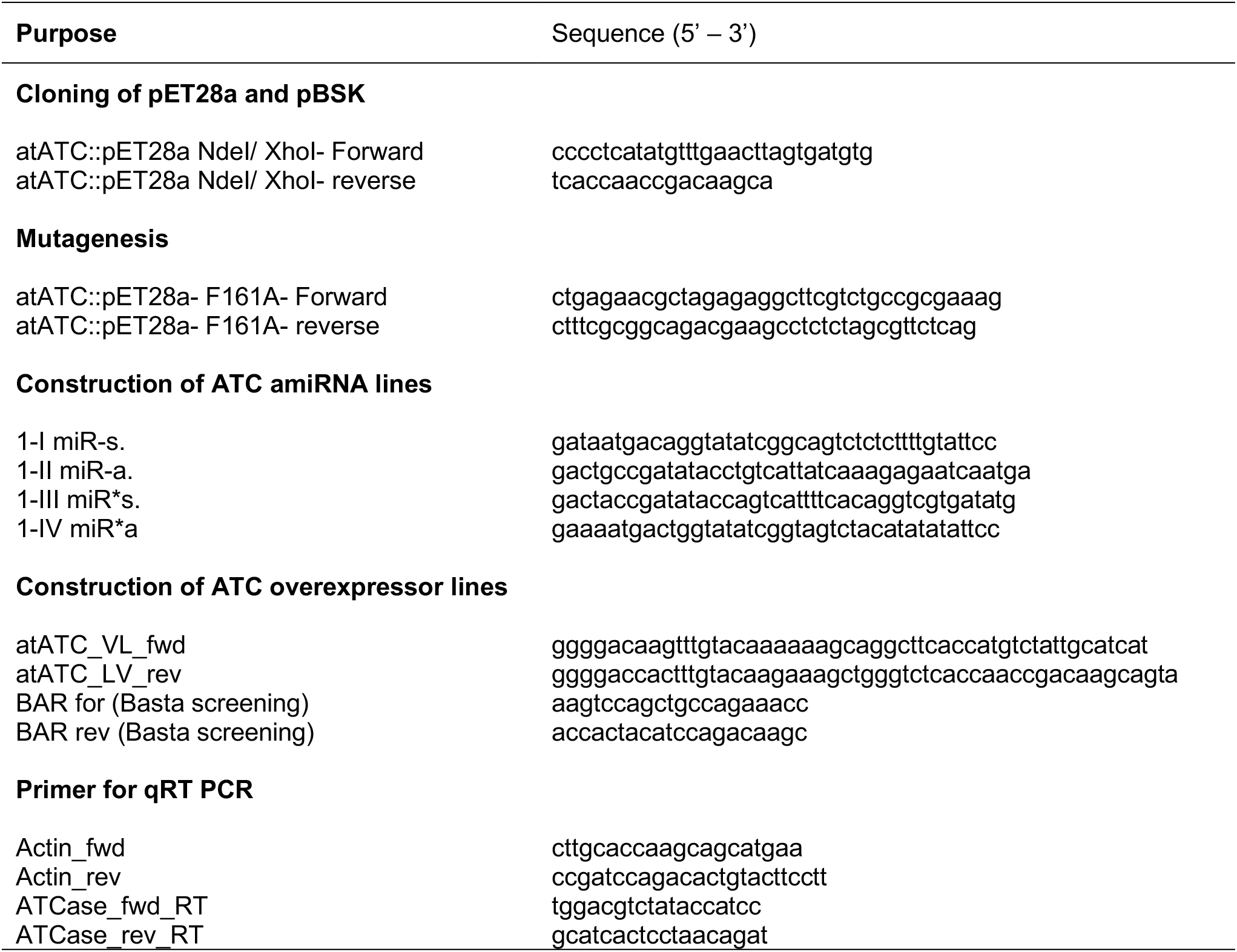
Primers used in this study.

---

**Supplementary Figure 1.**
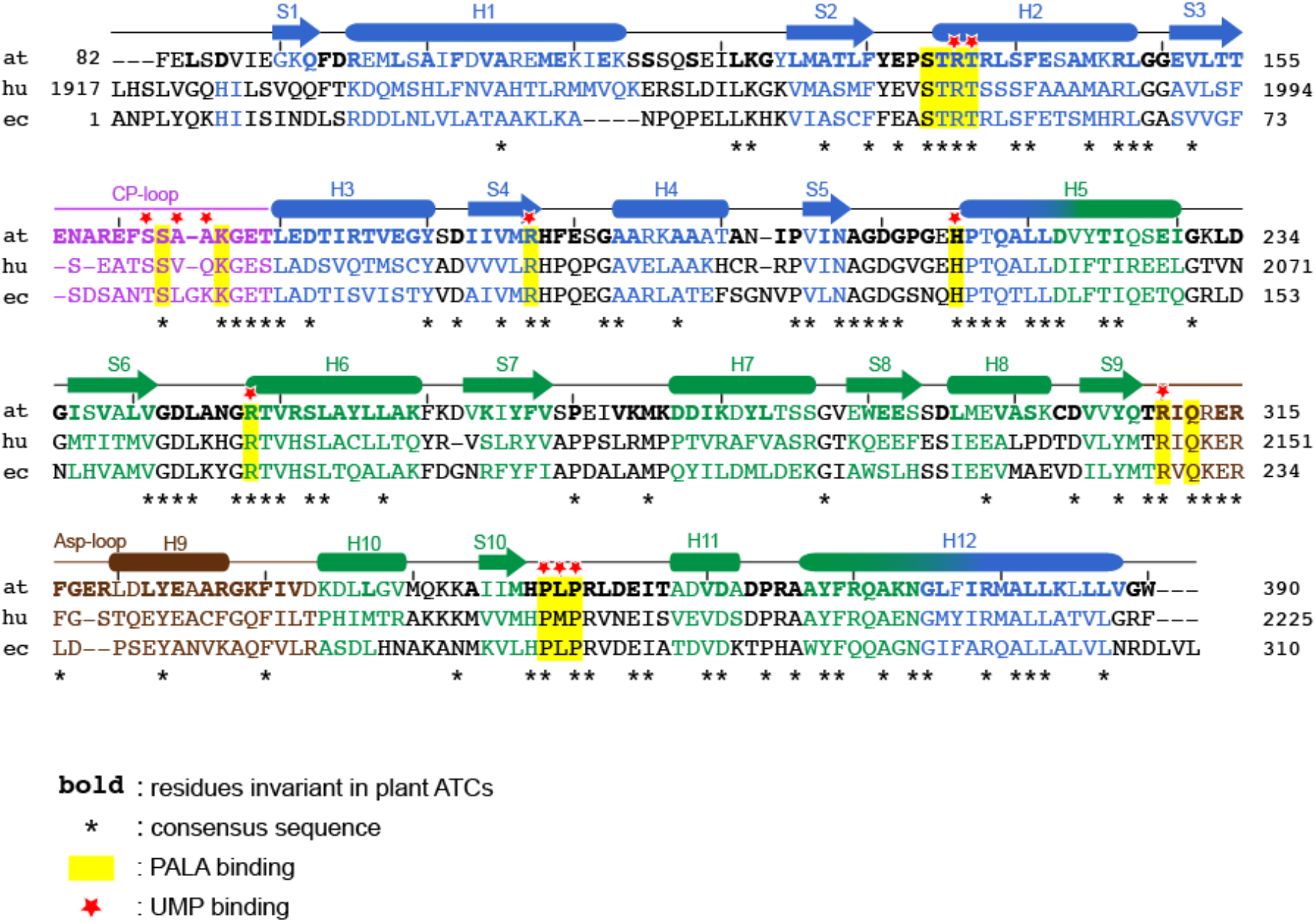
ATC sequence alignment. Structure-guided alignment of Arabidopsis (at), human (hu) and *E. coli* (ec) ATC sequences. The chloroplast transit peptide is not included. Asterisks indicate the consensus sequence. Sequence identity between Arabidopsis ATC and human or *E. coli* ATCs is 46% (138 of 302 aa, 97% covery) and 43% (131 of 302 aa, 97% covery), respectively. Conserved active site residues are shown on yellow background and residues interacting with UMP in atATC are indicated with a red star. Residues are colored according to the secondary structure, which is depicted above. α-Helices are shown as cylinders and β-strands as arrows. Residues in bold are found invariant in the following plant ATC sequences: *Pisum sativum* (Q43086 and Q43087), *Phaseolus vulgaris* (Phvul.008G270600.1), *Trifolium pratense* (Tp57577_TGAC_v2_mRNA1181 and Tp57577_TGAC_v2_mRNA2991), *Glycine max* (Glyma.02G293500.1, Glyma.14G021000.1 and Glyma.05G131400.1), *Oryza sativa* (Q9LD61), *Triticum aestivum* (partial sequence aa 122-390, TraesCS5A02G120200.1.; EnsemblPlants), *Camellia sinensis* (XP_028078246.1), *Eutrema salsungineum* (XP_006406396.1), *Capsella rubella* (XP_023641307.1), *Raphanus sativus* (XP_018490780.1), *Mircothlaspi erraticum* (CAA7053211.1), *Brassica oleracea* (XP_013638479.1), *Brassica napus* (XP_013728634.1), *Pistacia vera* (XP_031265500.1), *Citrus clementina* (XP_006427067.1), *Cajanus cajan* (XP_020237797.1), *Carica papaya* (XP_021906135.1), *Punica granatum* (XP_031407045.1), *Prunus dulcis* (BBH02036.1), *Hevea brasiliensis* (XP_021642880.1), *Vitis vinifera* (RVW30007.1), *Jatropha curcas* (XP_012070892.1), *Manihot esculenta* (XP_021593647.1), *Ricinus communis* (XP_002529466.2), *Prunus persica* (ONI13427.1), *Aquilegia coerulea* (PIA59089.1), *Striga asiatica* (GER28020.1), *Trema orientale* (PON91905.1), *Carpinus fangiana* (KAE8125493.1), *Cucurbita moschata* (XP_022932282.1) and *Aquilegia coerulea* (PIA59089.1).

**Supplementary Figure 2.**
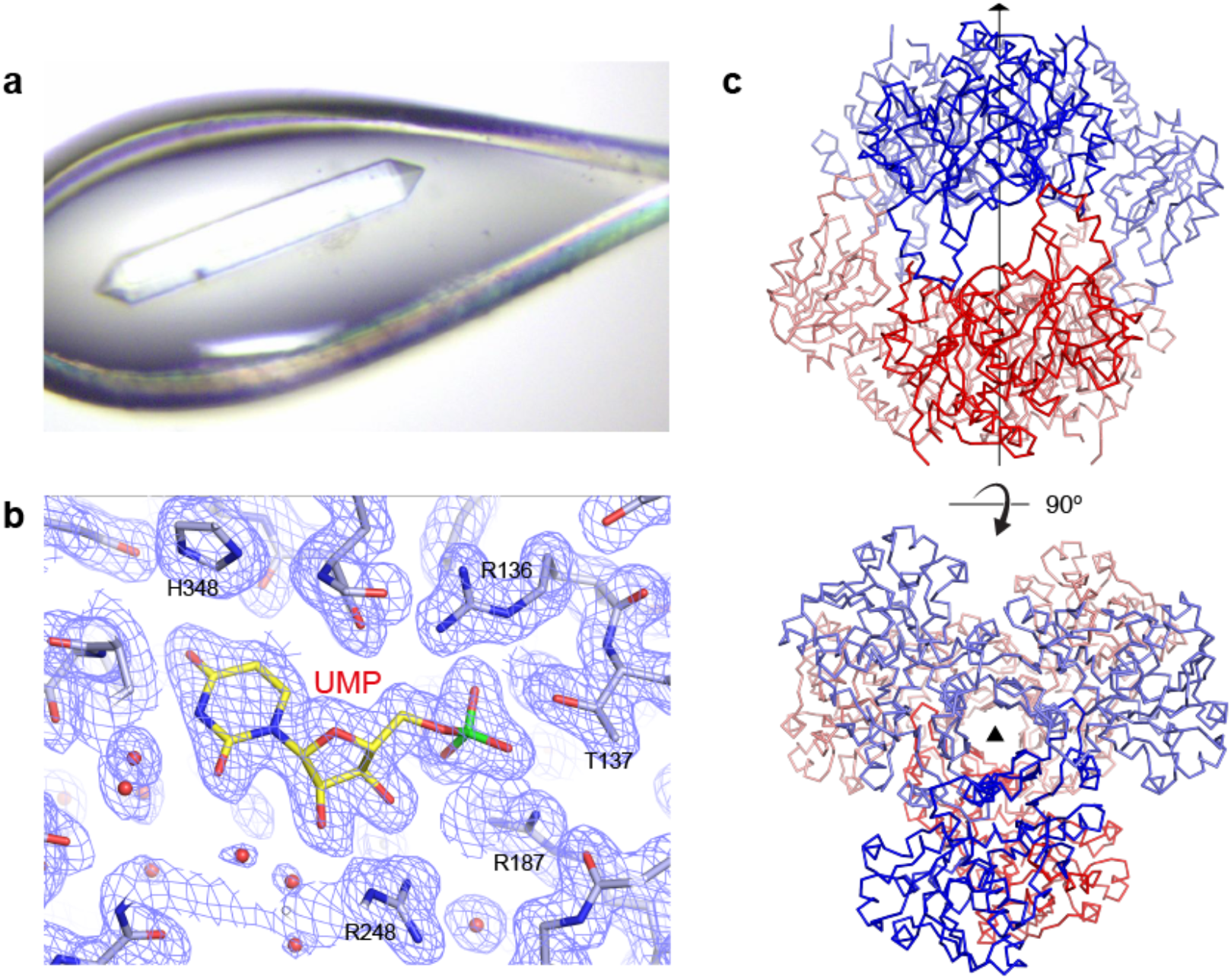
Structural determination of Arabidopsis ATC. **(a)** atATC crystal mounted on a cryo-loop during diffraction collection. **b)** Detail of the active site with UMP bound and 2F_obs_-F_calc_ electron density map represented in blue mesh. **c)** Perpendicular views of the two atATC subunits in the asymmetric unit colored in dark blue and red. Each subunit belongs to a different trimer that forms around the crystallographic three-fold axis. The other subunits in the trimers are depicted in lighter color.

**Supplementary Figure 3.**
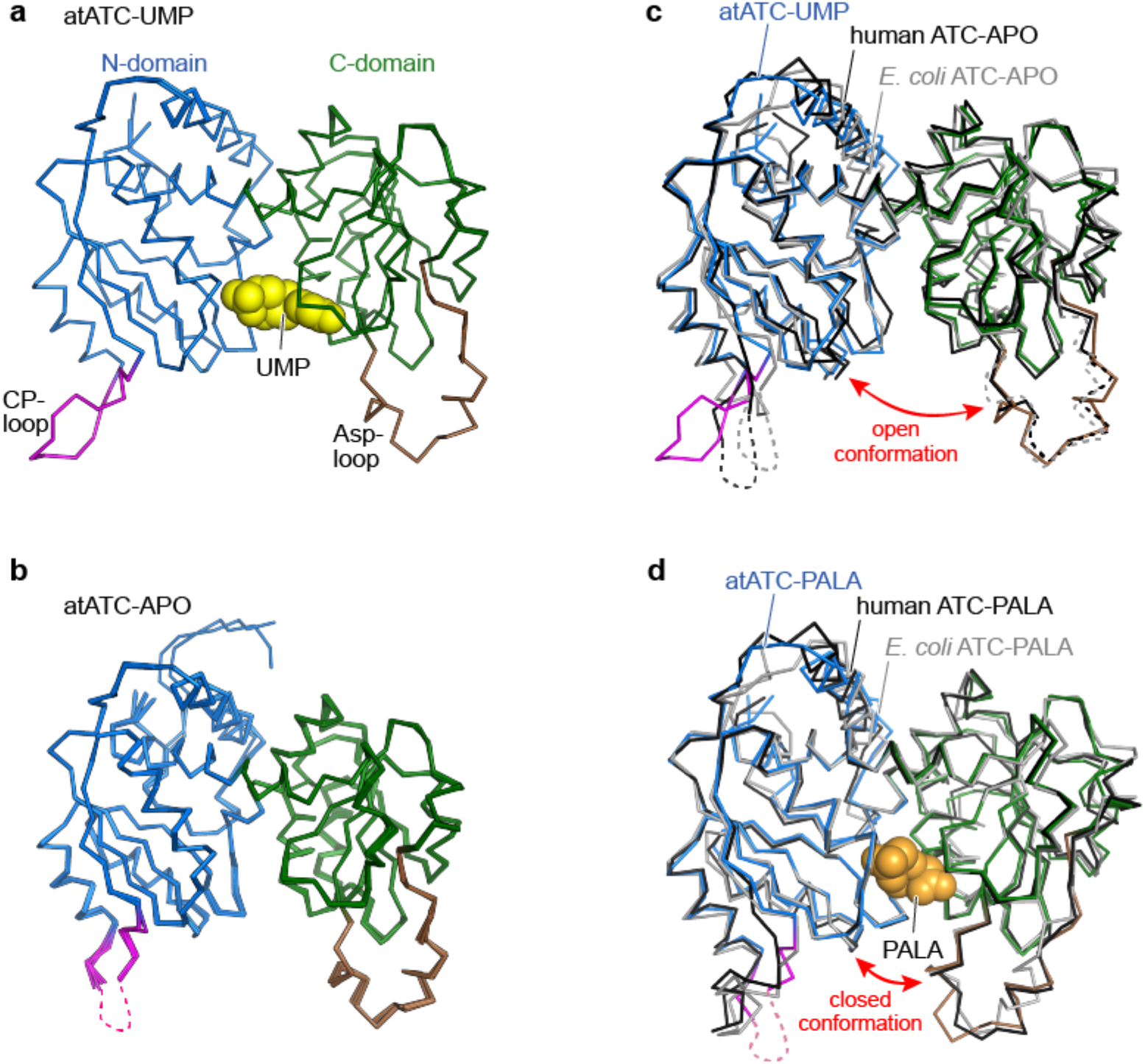
Comparison of atATC subunits within the different crystal structures. **a)** Superposition of the two protein subunits in the asymmetric unit of the atATC-UMP crystal. The entire polypeptide chains (311 Cα’s) superimpose with a rmsd= 0.24 Å. The subunits are represented in cartoon with the N- and C-domains depicted in blue and green, respectively. The CP-loop and the Asp-loop are colored pink and brown, respectively. UMP is shown as yellow spheres. **b)** Similar representation for the superposition of the six subunits in the atATC-APO crystal (rmsd=0.34-0.48 Å for 301-304 Cα’s). Dashed line indicates the flexibly disordered CP-loop. **c)** Superposition of atATC-UMP subunit with the apo structures of human and *E. coli* ATCs (PDB 5G1O and 3CSU). **d)** Superposition of atATC-PALA subunit with the PALA-bound subunits of human and *E. coli* ATCs (PDB 5G1N and 1D09).

**Supplementary Figure 4.**
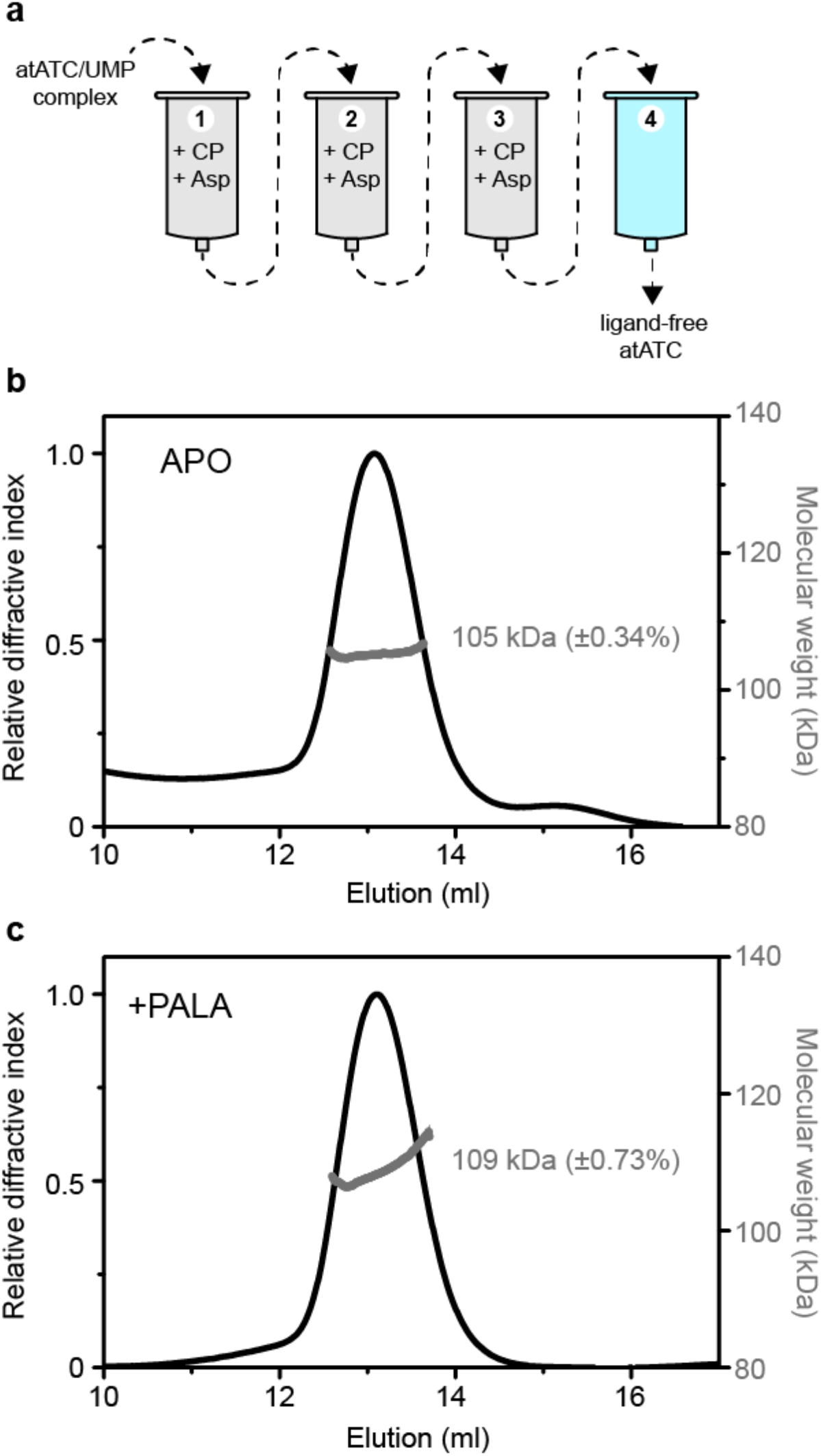
atATC free or bound to PALA remains as a homotrimer in solution. **a)** Scheme of the gel filtration procedure to remove UMP from atATC. The sample (2.5 ml at a concentration of 1 mg ml^−1^) is filtered through three consecutive 5 ml gel-filtration columns (PD-10 GE) equilibrated in a buffer solution containing 5 mM CP and 10 mM Asp. The sample elutes in 3.5 ml and is concentrated in an ultracentrifugation device to 2.5 ml before applying to the next column. In the last step, the sample is applied to a column equilibrated without substrates. The procedure lasts ~1 h and is performed at room temperature. **b,c)** SEC-MALS analysis of the UMP-freed sample **(b)** and pre-incubated with PALA **(c)**.

**Supplementary Figure 5.**
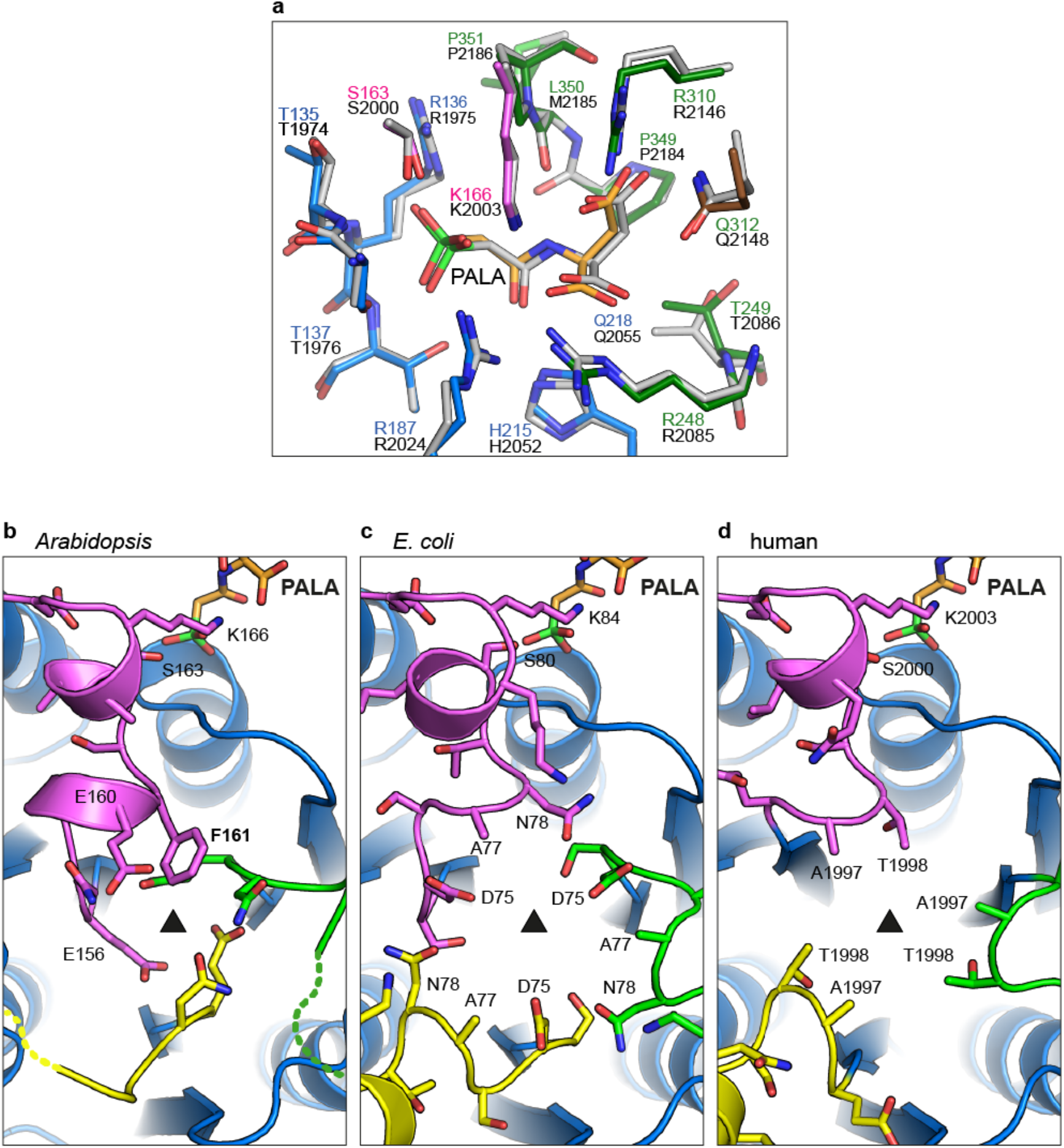
PALA binds to only one subunit per trimer. **a)** Superimposition of PALA-bound active sites of atATC and human ATC. atATC residues from the N-domain, the CP-loop or the C-domain are colored with C atoms in blue, pink or green, respectively, whereas PALA is depicted with C atoms in orange. Residue numbers are colored accordingly. The C atoms of human ATC are depicted light grey and residues are labeled in black. **b–d)** View along the threefold axis of Arabidopsis **(b)**, *E. coli* **(c)** and human **(d)** ATCs bound to PALA. The CP-loops from the three subunits in the trimer are shown in different colors.

